# DIAproteomics: A multi-functional data analysis pipeline for data-independent-acquisition proteomics and peptidomics

**DOI:** 10.1101/2020.12.08.415844

**Authors:** Leon Bichmann, Shubham Gupta, George Rosenberger, Leon Kuchenbecker, Timo Sachsenberg, Oliver Alka, Julianus Pfeuffer, Oliver Kohlbacher, Hannes Röst

**Affiliations:** Department of Computer Science, Applied Bioinformatics, University of Tübingen, Germany; Institute for Cell Biology, Department of Immunology, University of Tübingen, Germany; Donnelly Center for Biomolecular research, University of Toronto, Toronto, Canada; Department of Systems Biology, Columbia University, New York, NY, USA; Institute for Informatics, Freie Universität Berlin, Berlin, Germany; Zuse Institute Berlin, Berlin, Germany; Institute for Biomedical Informatics, University of Tübingen, Germany; Institute for Translational Bioinformatics, University Hospital Tübingen, Germany

## Abstract

Data-independent acquisition (DIA) is becoming a leading analysis method in biomedical mass spectrometry. Main advantages include greater reproducibility, sensitivity and dynamic range compared to data-dependent acquisition (DDA). However, data analysis is complex and often requires expert knowledge when dealing with large-scale data sets. Here we present DIAproteomics a multi-functional, automated high-throughput pipeline implemented in Nextflow that allows to easily process proteomics and peptidomics DIA datasets on diverse compute infrastructures. Central components are well-established tools such as the OpenSwathWorkflow for DIA spectral library search and PyProphet for false discovery rate assessment. In addition, it provides options to generate spectral libraries from existing DDA data and carry out retention time and chromatogram alignment. The output includes annotated tables and diagnostic visualizations from statistical post-processing and computation of fold-changes across pairwise conditions, predefined in an experimental design. DIAproteomics is open-source software and available under a permissive license to the scientific community at https://www.openms.de/diaproteomics/.

## INTRODUCTION

Recently, data-independent acquisition (DIA) using sequential windowed acquisition of all theoretical fragment ion mass spectra (SWATH-MS)^1^ has attracted much attention in the field of proteomics due to its ability to overcome shortcomings of the classical data-dependent (DDA) strategy.^2–6^ Because of its outstanding performance in reproducibility and quantification, DIA is likely to become the state-of-the-art technology in clinical mass spectrometry (MS).^7^ In addition, recent tailored applications of DIA have enabled new approaches for the chemoproteomic screening of drug targets.^8^ The main advantages are its capacity to (1) acquire fragment spectra in a reproducible grid-based fashion over the entire mass and retention time range, (2) sample fragment spectra for nearly all precursor ions present in a sample, and (3) enable to trace elution profiles of fragments and integrate their quantities at a greater dynamic range.^9^ Yet, this comes at the cost of increased complexity of the acquired mass spectra, due to simultaneous fragmentation of multiple precursor ions, which requires appropriate methods for spectra identification.^10^ Nonetheless, DIA has the promising potential to achieve a greater identification rate and quantification range, higher reproducibility, and fewer missing values than DDA.

A key step to process DIA data is the generation of high-quality spectral libraries to identify the complex DIA spectra with higher sensitivity.^11^ These spectral libraries can be derived from previously acquired DDA measurements by selectively annotating and storing peak intensities and other properties from confident peptide spectrum matches across multiple samples. Public repositories such as PRIDE^12^, the PeptideAtlas Project^13^, the SWATHAtlas^14^ or the SysteMHCAtlas^15^ provide collections of aggregated spectral libraries from large DDA datasets such as the human proteome or spectral libraries of other species or specific contexts.^16^ Alternatively, recently developed *in silico* methods that utilize advanced machine learning strategies to predict peptide fragment intensities can be applied.^17–20^ However, the library should match the settings of instrument and acquisition method to which the respective DIA experiment will be compared to, as different instruments, ionization methods, and corresponding parameters such as collision energies produce vastly different fragment spectra patterns. Finally, as an additional alternative, library free approaches for the deconvolution of DIA data have been proposed to overcome the limitations and dependencies of spectral libraries.^10^

With increasing amounts of MS measurements recorded in both DDA and DIA acquisition mode deposited in publicly available data repositories^12^, there is a need for automated high-throughput data analysis pipelines. As the parametrization of DIA search algorithms and the choice of a spectral library can strongly influence the analysis results, flexible and scalable software solutions for high-performance computing systems are required to provide ways to efficiently reprocess and compare analysis results using large amounts of existing data. This includes the automated generation of spectral libraries from available DDA measurements and the alignment of their transition retention times into the same space. Previously, multiple different software solutions have been applied to process large-scale DIA data^21–24^, however, their application often requires expert-knowledge and a combination of several post-processing procedures or manual interaction at various analysis steps.

We address this gap in available software solutions by introducing DIAproteomics, a versatile, high-throughput analysis pipeline for DIA proteomics and peptidomics MS measurements. It achieves a high degree of automation and scalability from single users to large high-performance computing (HPC) environments, by integrating well-established tools such as the OpenSwathWorkflow^22^ for DIA library search, provided through the OpenMS software toolbox for computational mass spectrometry^25,26^. The false discovery rate (FDR) is estimated by the PyProphet algorithm^27^, followed by chromatogram alignment as a post-processing step using the DIAlignR software^28^. Moreover, it provides the option to use it either by specifying a particular existing spectral library and retention time standards or by generating the spectral library and selecting suitable pseudo-iRTs from existing DDA measurements and search results. Ultimately, statistical post-processing provided through MSstats^29^ ensures reliable analysis results.

DIAproteomics is containerized and implemented using the workflow language Nextflow^30^, leveraging the capabilities of the powerful Nextflow execution engine to seamlessly run on single desktop computers and scale up to large-scale HPC or cloud environments. As part of the nf-core repository for reproducible bioinformatics workflows^31^ it adheres to the corresponding strict standards. Ultimately, a browser-based user interface accompanies the workflow and allows easy-to-use parametrization and execution.

## METHODS

### Pipeline architecture

DIAproteomics is an automated analysis pipeline that can be broadly partitioned into the following parts: Optional spectral library and iRT generation from provided DDA data, optional spectral library merging and RT alignment, DIA library search, false discovery rate (FDR) estimation, MS2 chromatogram alignment across runs, and output summarization (Figure 1). Each of these parts involves one or more required or optional steps within the workflow (Supplementary Information Table S1 and Figure S1). An experimental design needs to be provided in the form of an input sample sheet specifying DDA and DIA samples, libraries or iRT standards that should be coprocessed in one batch.

**Figure 1:**
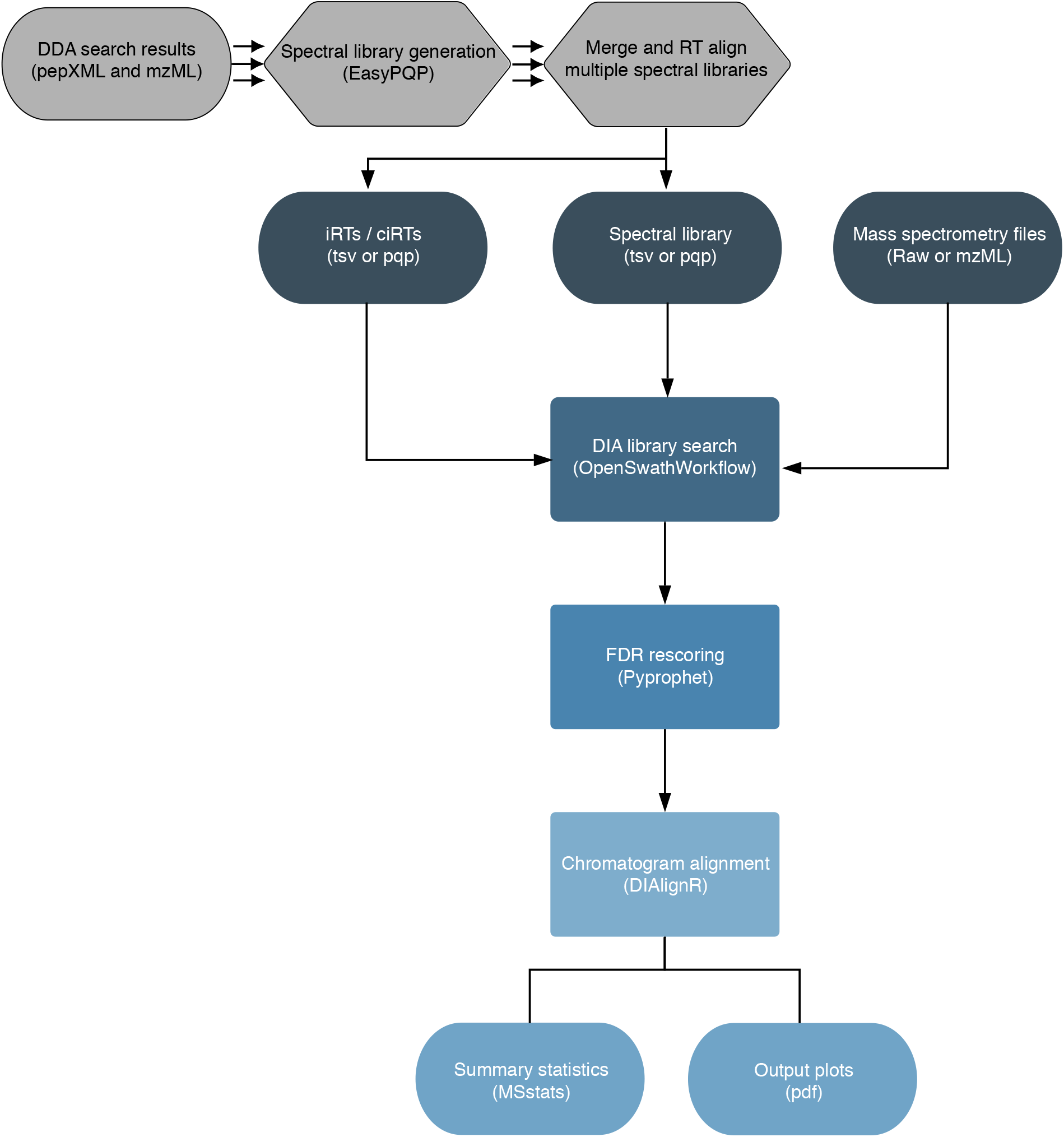
Simplified scheme of the DIAproteomics workflow. The input to the pipeline can be either spectral libraries and iRTs generated and combined from DDA raw data (optional, in gray) or otherwise an existing spectral library and internal retention time standards (iRTs). Next, targeted extraction is performed by searching the DIA-SWATH MS raw files with the spectral library using the OpenSwathWorkflow. The false discovery rate (FDR) is assessed applying PyProphet subsequently. Next, chromatograms are aligned using the DIAlignR software. Finally, the output is statistically post-processed with MSstats and visualized.

#### Spectral library generation

In a first, optional step provided DDA raw MS measurements (Thermo Raw vendor format) are converted to the open, XML-based mzML format^32^. Next, the library is generated using EasyPQP (available at https://github.com/grosenberger/easypqp) which matches the provided search results (for example in pepXML format) and the corresponding DDA raw measurements to annotate and store peptide transitions and their properties in a tab-separated table^33^. The library is transformed into an assay containing a specified number of transitions of band y-ions falling into a custom mass-to-charge range. Subsequently, decoy transitions that can be generated by OpenMS in multiple ways such as reversed or shuffled are added to the library. Finally, the generated library will be exported in the peptide query parameter (pqp) sqlite-based data format. Optionally, all steps of the library and decoy generation can be skipped, and an existing library can be used instead.

#### Pseudo iRT generation

If specified, a given number of highly confident peptide identifications spanning the entire RT range will be selected and exported to serve as iRT standards in the DIA library search step. This is important, for example, if no iRT standard kit was spiked into the samples before the DIA measurements. Selected iRTs will be exported in the peptide query parameter (pqp) sqlite-based data format. However, if provided, a set of user-defined iRTs can be used instead.

#### Spectral library merging

If multiple libraries per sample are provided, for example when stemming from a set of technical replicates, the libraries can be optionally merged and will then undergo a linear RT alignment onto the same reference. When merging is enabled, the best scoring peptide identification is kept in the library omitting a lower scoring duplicate.

#### Spectral library RT alignment

When RT alignment is enabled, the multiple input spectral libraries will be pairwise aligned onto the same reference. This is achieved by computing a minimum spanning tree connecting all provided libraries by shared peptide overlap. (Supplementary Information Figure S3) Hence the library having the highest overlap in shared peptides with all other libraries will be the central reference for the other libraries. Importantly, this strategy is also applicable when aligning very distant libraries onto the same reference that share no consensus peptide identifications among all libraries.^34^ However, it requires peptides to be shared between all pairs of libraries, resulting in a connected tree.

#### DIA spectral library search

In a first, optional step provided DIA raw MS measurements (Thermo RAW vendor format) may be converted to the mzML XML-based format. Next, DIA library search is carried out using the OpenSwathWorkflow, implemented within the OpenMS toolbox. The spectral library and iRT standards are used to search all input DIA raw measurements individually with a customizable parametrization. The swath windows can be determined from the data. Finally, extracted ion chromatograms (XICs) of the searched peptide transitions (mzML) are exported and the output features and transition properties are stored in OpenSwathWorkflow files (osw).

#### False discovery rate estimation

The OpenSwathWorkflow output files (osw) are merged samplewise as defined in the experimental design (sample sheet). The merged file is then scored using the PyProphet target-decoy FDR estimation procedure. Finally, the level of confidence such as local transition- or global peptide or protein level-based can be defined.^27^ The PyProphet scoring results will then be exported as a tab-separated table per DIA MS run and the results will be visualized in a pdf report.

#### MS2 chromatogram alignment

As the last processing step, the extracted and scored MS2 chromatograms will be aligned using the DIAlignR software. This involves matching chromatograms between runs that can be aligned and integrating their transition areas. The sum of the integrated areas per peptide will be reported as peptide quantities in a TSV file. For this procedure, DIAlignR provides several FDR estimates that can be customized within the workflow to define cut-offs for transitions that should be excluded from matching between runs.^28,35^

#### Output summarization

The output is summarized in a pairwise manner on peptide or protein level using the MSstats post-processing software^29^. In addition, it is possible to export a number of diagnostic plots illustrating peptide and protein identification results, their quantities and properties.

### Implementation

The DIAproteomics pipeline is implemented in the Nextflow workflow programming language^30^, based on the nf-core community template for reproducible bioinformatics workflows^31^. Support for multiple functionalities is provided such as for various container systems (e.g., docker, singularity, podman), environment management platforms (e.g. Conda), the user interface and support for the execution on high-performance computing systems such as google cloud or amazon web services. Each step of the workflow is executed as an independent process allowing efficient, parallel processing of large amounts of data.

Most of the inner functions and the file format handling is provided through the OpenMS v.2.5.0 toolbox for computational mass spectrometry^26^. Specifically, this includes the handling of spectral libraries, assay and decoy generation and the implementation of the OpenSwathWorkflow^22^. Spectral library generation from DDA data is carried out by EasyPQP v.0.1.7. A customized python v.3 script is executed to merge multiple libraries and compute the minimum spanning tree for RT alignment using the module NetworkX v.2.4^36^. False discovery rate estimation on merged OpenSwathWorkflow^22^ output files is achieved using functionalities of PyProphet v.2.1.4^27^, MS2 chromatogram alignment and integration of peptide quantities is achieved using the ‘alignTargetedRuns’ function of the DIAlignR software 1.2.0^28^. The ‘groupComparison’ function using ‘highQuality’ feature subsets within MSstats v. 3.20.1^29^ is carried out to compute protein level statistics and pairwise comparisons of protein fold-changes and significance across conditions. Finally, output visualizations are created using the R software libraries gplots and ggplot2.

### Parametrization

The DIAproteomics workflow is highly flexible and each execution step provides various parameters that can be customized for specific instrumental and experimental settings. An overview over available parameters and a short description is provided at https://nf-co.re/diaproteomics. The default parametrization has been benchmarked multiple times in the past^23,37^. It involves spectral library assay generation with the six most intense b- and y-ion transitions falling into the precursor mass range of 400 to 1200 m/z and a fragment mass range of 350 to 2000 m/z. The default setting for decoy transition generation is shuffling. The extraction of MS1 precursor and MS2 fragment transitions is carried out using a mass extraction window of 10 and 30 ppm respectively and an RT extraction window of 600 seconds for the targeted extraction of the OpenSwathWorkflow. The false discovery rate estimation is performed on global protein level involving an LDA based target-decoy separation. The MS2 chromatogram alignment requires transitions to satisfy several FDR thresholds. For global alignment, high quality peaks are selected with globalAlignmentFdr (set to 0.01) cutoff. A peak will only be matched across runs if at least one run has estimated FDR is below of 0.01 (UnalignedFDR). It will then be compared to matching peaks in other runs below a higher maximum FDR threshold of 0.05 (MaxQueryFDR). This is an advantage over common strategies used in DDA to allow matching between runs, since no FDR cut-off can be set for these approaches.

### Reanalysis of publicly available data sets

A concise benchmark on the publicly available multi-center benchmark study data set by Navarro et al^23^ (PRIDE PXD002952) was carried out using a Human, *E. coli* and Yeast mixture HYE124. The TripleToF 6600 and 64 variable swath window instrument setting was chosen applying the default parametrization of DIAproteomic v1.1.0, adjusting the precursor and fragment mass tolerances to 50 and 30 ppm respectively. SCIEX wiff files were converted to mzML using the proteowizard msconvert software. Converted mzML files were further centroided on both MS levels using the OpenMS tool PeakPickerHiRes v.2.5.0.

Ultimately, to ensure the capability of the DIAproteomics pipeline v1.1.0 to process publicly available proteomics data sets, several HeLa cell line Thermo orbitrap high resolution MS runs from the PRIDE project PXD003179^38^ were reanalyzed using the default settings. The procedure was automated and integrated as a continuous integration full size test on Amazon web services (AWS) that can be actively run to verify the pipeline’s functionality.

## RESULTS AND DISCUSSION

### DIAproteomics facilitates the analysis of large-scale DIA-SWATH MS measurements

DIAproteomics is a versatile analysis pipeline for processing of large-scale proteomics and peptidomics DIA-SWATH mass spectrometry runs. As its implementation is based on the nf-core template for reproducible bioinformatics workflows, DIAproteomics provides a web-based browser interface that can be customized. It allows to get an overview and adjust the available parameters grouped into several categories and documenting their functions in short to longer expandable descriptions. Several sample sheets that annotate batch identifiers and conditions to each sample as defined by the experimental design serve as input to the pipeline. (Figure 2)

**Figure 2:**
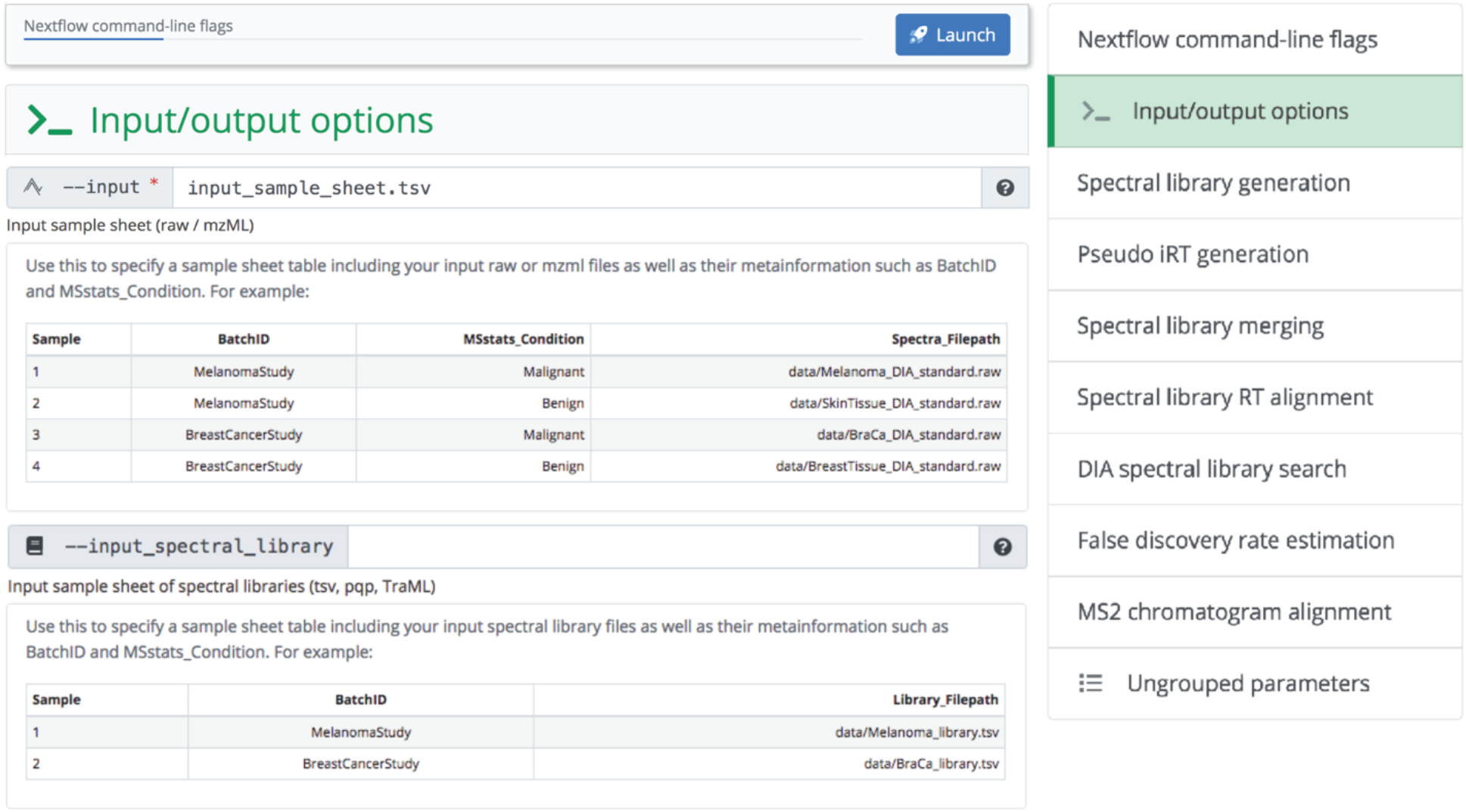
Input / output options as available through the nf-core provided user-interface. Spreadsheets serve as input to the pipeline defining the experimental design of raw files, spectral libraries and their corresponding conditions and batch identifiers (BatchID). Upon submission of the job MS runs are grouped by their BatchID and coprocessed.

Whenever possible each step of the pipeline is executed and submitted individually for processing by the computing infrastructure. In this way, the processing of multiple large batches of files can be efficiently parallelized. On the other hand, if steps allow to combine multiple files, the workflow groups the files according to the experimental design and co-processes them. This occurs for instance when merging and aligning multiple spectral libraries or when carrying out a global FDR estimation on merged DIA search results.

Depending on how the parameters were set within the major categories, the input and output files may vary. Most importantly, it can be defined whether one or multiple existing spectral libraries should be used or whether the spectral libraries should be generated from matching DDA raw files and peptide identification results. (Figure 3) Yet, many more settings for each of the parameter categories are available and can be tailored to specific problem settings and MS instrument requirements.

**Figure 3:**
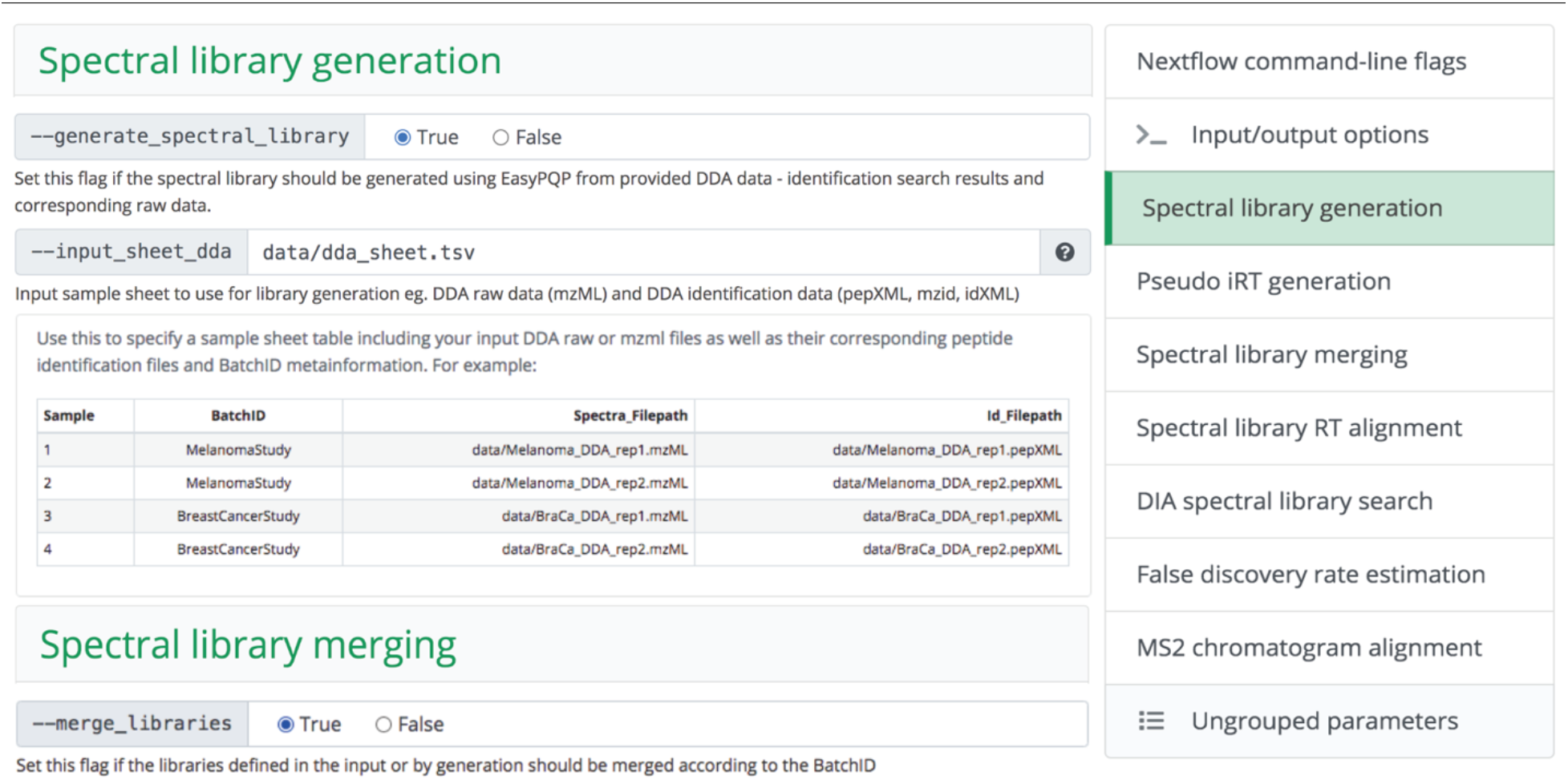
(Optional) Generation of spectral libraries from DDA raw data as it can be defined through the nf-core provided user-interface. A spreadsheet annotates DDA raw files, corresponding peptide identification results and their batch identifiers (BatchID). If specified, multiple spectral libraries from several MS runs of the same batch will be merged upon submission of the job.

### Statistical post-processing and diagnostic output visualization

The output of the DIAproteomics pipeline is by default a set of tables as well as illustrations summarizing peptide or protein amount and quantities and scoring results. Moreover, important intermediate results such as the generated libraries, the output of the DIA spectral library search, and XICs are reported. Most importantly, the detailed target-decoy score distribution results and their visualizations as exported from PyProphet are deposited in the output directory. The MSstats post-processing software is run on the determined peptide or protein quantities. This results in the statistically sound estimation of pairwise fold changes and their significance across the conditions defined in the experimental design that are as well visualized in comparative plots such as a Volcano visualization. In addition, more diagnostic plots can be generated listing the number of peptides and proteins identified, their properties such as the charge distribution, RT deviation between the spectral library and DIA measurement to assess the performance of the iRT alignment (Figure 4). Finally, if specified a heatmap of peptide quantities and missing values across all DIA MS runs is exported.

**Figure 4:**
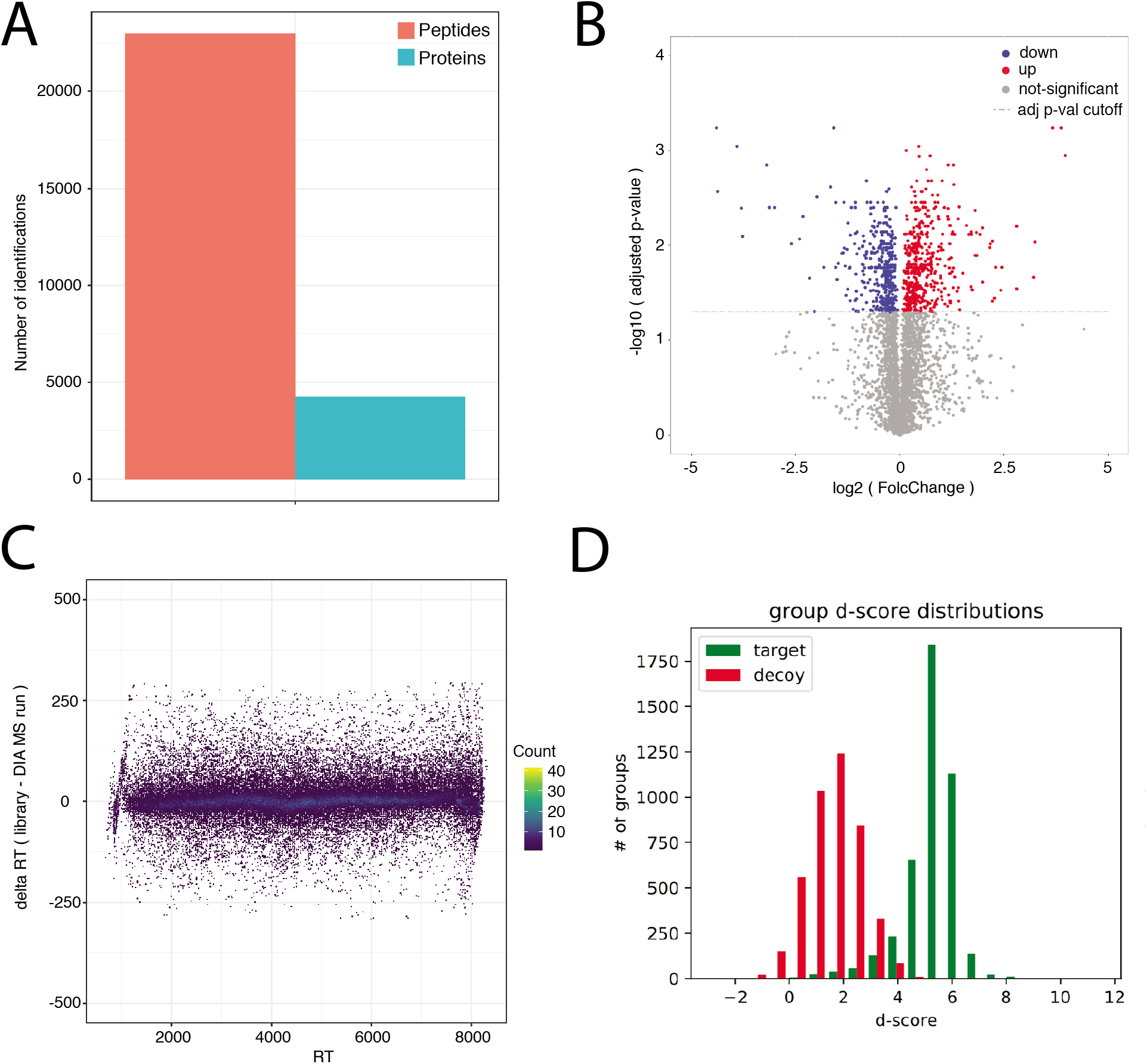
Several diagnostic visualizations of the DIAproteomics output can be generated. The results of reprocessing of the publicly available dataset PRIDE-PXD003179 are shown here as an example. A) Peptide and protein identification counts. B) Volcano plot of differentially regulated proteins (red up, blue down) proteins across conditions. C) Deviation of Spectral library and MS run in retention time (RT) over the entire RT range. D) Target and decoy d-score distribution as computed by PyProphet to assess the false discovery rate (FDR).

Finally, quantification performance of the DIAproteomics was additionally compared to the results of a multi-center benchmark study.^23^ As a result we were able to reproduce the log-fold changes of the used human, E. coli and yeast mixture at defined ratios. (Supporting Information Figure S2)

### Run time considerations

The runtime of the DIAproteomics workflow depends on its parametrization. For example, if spectral library generation from DDA data is chosen and the number of samples and batches that are analyzed in one submission. We assessed the computational runtime and required resources making use of Amazon web services (AWS) cloud infrastructure and the German network for bioinformatics infrastructure (de.NBI) cloud HPC node with 28 cores and 64 GB. The analysis of six DIA-SWATH MS runs applying a library and pseudo iRTs generated from three DDA MS runs was carried out in approximately 2h and 10min using AWS. (Figure 5)

**Figure 5:**
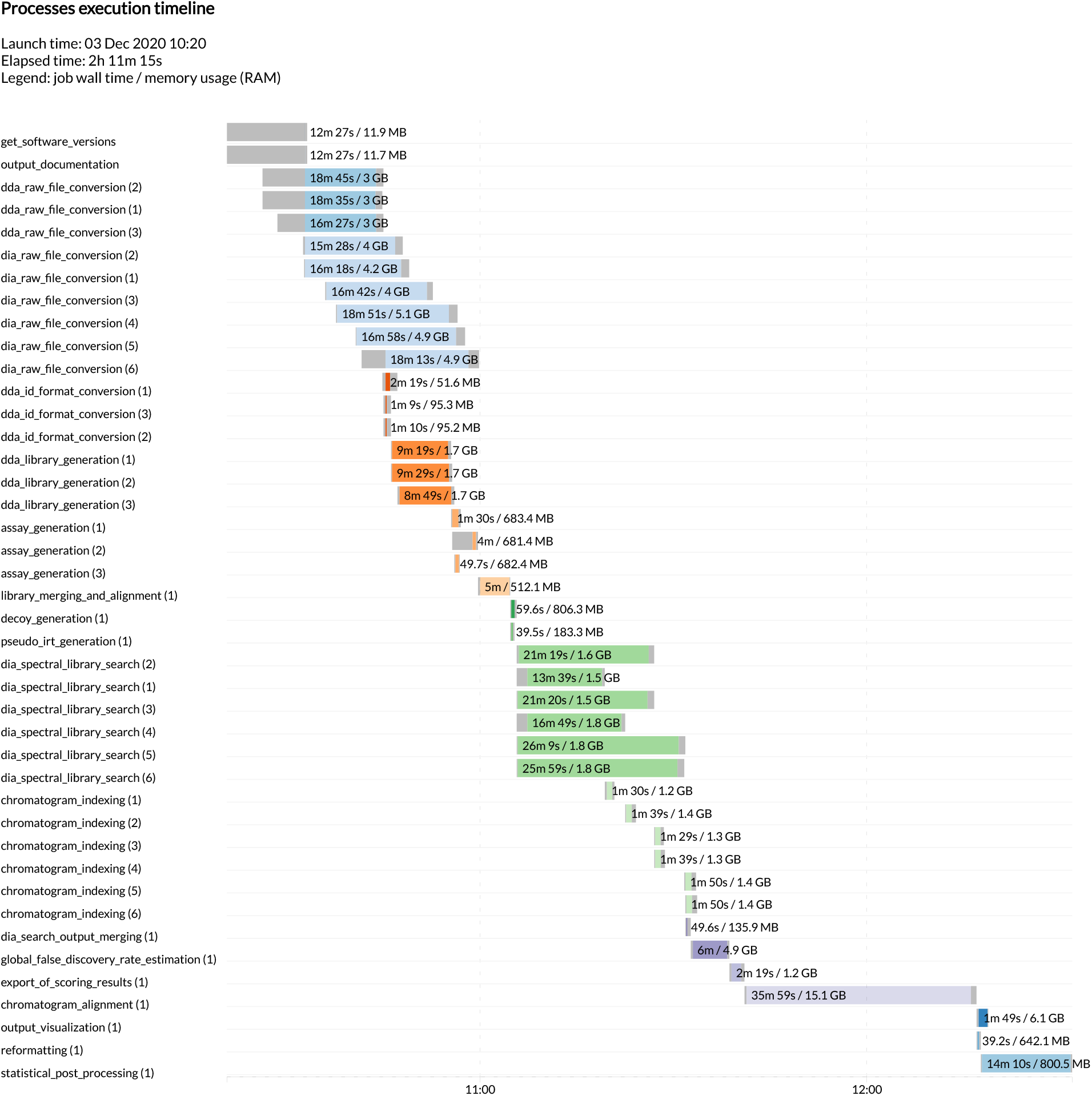
Detailed overview on run times and memory usage of all integrated steps of the DIAproteomics pipeline when processing the PRIDE dataset PXD003179 on the Amazon webservice cloud infrastructure.

## CONCLUSION

In this work we present DIAproteomics, a flexible computational workflow to automatically process large scale DIA-SWATH MS based proteomics and peptidomics studies on diverse computational systems. It combines all steps including the optional generation of spectral libraries from DDA data and the essential DIA library search, FDR estimation and chromatogram alignment. Implementation and sharing the workflow as part of the nf-core initiative for reproducible bioinformatics research provides an easy-to-use user interface as well as reproducible, well tested analysis. The DIAproteomics pipeline is provided for free to the science community, with the purpose to enable easier access, as well as automated and reproducible analysis of DIA-SWATH MS based proteome research.

## ASSOCIATED CONTENT

The workflow is freely available under an open-source license as Nextflow implementation in the nf-core bioinformatics workflow repository: https://www.openms.de/diaproteomics/. Moreover, a detailed documentation regarding parameters and pipeline output can be found at: https://nf-co.re/diaproteomics

Supporting Information Available:

Supporting Material Table S1. Details on all steps in the Nextflow workflow implementation

Supporting Material Figure S1. Command line execution report of the workflow

Supporting Material Figure S2. Benchmarking quantification performance

Supporting Material Figure S3. Pairwise RT alignment option for merging multiple spectral libraries

## Author Contributions

L.B. constructed the pipeline, carried out the data analysis and wrote the paper. S.G. created DIAlignR and G.R. created EasyPQP – two software tools that are essential components of the pipeline and both supported their integration and debugging into the workflow. L.K., T.S., J.P., O.A. assisted in reviewing the source code and suggested architecture and parameter changes. O.K. and H.R. were involved in the study design. All authors discussed and commented on the manuscript.

## ACKNOWLEDGMENT

We would like to thank all nf-core and OpenMS team members, in particular Phil Ewels and Gisela Gabernet for supporting the development and debugging of the pipeline as well as for the provision of the template. In addition, we would like to thank the Quantitative Biology Center in Tübingen (QBiC) for hosting a productive software developer meeting.

## ABBREVIATIONS

LC-MS/MS: liquid chromatography coupled mass spectrometry;
MS: mass spectrometry;
DIA: data-independent acquisition;
DDA: data-dependent acquisition;
FDR: false discovery rate;
XIC: extracted ion chromatogram;
HPC: high-performance computing

