## Supplemental Information for "DIAproteomics: A multi-functional data analysis pipeline for data-independent-acquisition proteomics and peptidomics"

Supporting Information Available:

Supporting Material Table S1. Details on all steps in the nextflow workflow implementation
Supporting Supporting Material Figure S1. Command line execution report of the workflow
Supporting Supporting Material Figure S2. Benchmarking quantification performance
Supporting Material Figure S3. Pairwise RT alignment option for merging multiple spectral libraries

Table S1 – Details on all steps in the nextflow workflow implementation

| Step | Requirement | Name | Description | |
|---|---|---|---|---|
| **1** | (optional) | DDA raw file conversion | DDA input files are converted into mzML from Thermo Raw vendor format. | Spectral library and iRT generation, merging and RT alignment |
| **2** | (optional) | DDA library generation | Spectral libraries are generated from DDA raw files and search results using EasyPQP. | |
| **3** | (optional) | Assay generation | Spectral library transitions are filtered by specific settings. | |
| **4** | (optional) | Library merging and alignment | If specified multiple generated libraries will concatenated and undergo linear RT alignment. | |
| **5** | (optional) | Pseudo iRT generation | A specified number of high-confidence peptide identifications are selected for the use as iRTs. | |
| **6** | (optional) | Decoy generation | Decoy transitions are appended to the input library. | |
| **7** | (optional) | DIA raw file conversion | DIA input files are converted into mzML from Thermo Raw vendor format. | DIA library search |
| **8** | required | DIA spectral library search | The OpenSwathWorkflow searches the DIA input files with either the provided or generated spectral library and iRTs. | |
| **9** | required | DIA search output merging | The OpenSwathWorkflow output results of the provided DIA measurements is merged together | FDR estimation |
| **10** | required | Global false discovery rate estimation | The OpenSwathWorkflow target-decoy output results are rescored on global peptide or protein level. | |
| **11** | required | Export of scoring results | The PyProphet scoring output results are exported per MS run and a report visualizing target and decoy scores is created. | |
| **12** | required | Chromatogram indexing | Extracted chromatograms exported from the OpenSwathWorkflow are reindexed.. | |
| **13** | required | Chromatogram alignment | Extracted chromatograms are realigned, matched between runs and quantified by the DIAlignR software. | Chromatogram alignment |
| **14** | (optional) | Reformatting | The DIAlignR output is reformatted to be passed onto the MSStats software. | |

| | | | | |
|---|---|---|---|---|
| **15** | (optional) | Statistical post processing | Analysis carried out by the MSStats software aggregating results on protein or peptide level. | |
| **16** | required | Output visualization | Several diagnostic plots are generated summarizing the output. | Output summary |

Figure S1. Command line execution report of the workflow

```
bash-4.2$ ./nextflow run diaproteomics -profile singularity,test_full
N E X T F L O W  ~  version 20.07.1
Launching `diaproteomics/main.nf` [insane_plateau] - revision: bda910a6f9
----------------------------------------------------
                                        ,--./,-.
        ___     __   __   __   ___     /,-._.--~'
  |\ | |__  __ /  ` /  \ |__) |__         }  {
  | \| |       \__, \__/ |  \ |___     \`-._,-`-,
                                        `._,._,'
  nf-core/diaproteomics v1.0dev
----------------------------------------------------
Run Name            : insane_plateau
Spectral Library    : generate spectral library from DDA data
Max Resources       : 16 GB memory, 32 cpus, 2d time per job
Container           : singularity - nfcore/diaproteomics:dev
Output dir          : ./results
Launch dir          : /nfs/wsi/abi/scratch/leonb/DIAproteomics
Working dir         : /nfs/wsi/abi/scratch/leonb/DIAproteomics/work
Script dir          : /nfs/wsi/abi/scratch/leonb/DIAproteomics/diaproteomics
User                : bichmann
Config Profile      : singularity,test_full
Config Profile Description: Full test dataset to check pipeline function
Config Files        : /nfs/wsi/abi/scratch/leonb/DIAproteomics/diaproteomics/nextflow.config
----------------------------------------------------
executor >  local (42)
[b9/9eec08] process > get_software_versions                    [100%] 1 of 1 ✔
[ab/ac3349] process > dda_raw_file_conversion (1)              [100%] 3 of 3 ✔
[f7/135ec1] process > dda_id_format_conversion (1)             [100%] 3 of 3 ✔
[b9/20b209] process > dda_library_generation (3)               [100%] 3 of 3 ✔
[6b/379821] process > assay_generation (3)                     [100%] 3 of 3 ✔
[2c/61ed25] process > library_merging_and_alignment (1)        [100%] 1 of 1 ✔
[be/25b263] process > pseudo_irt_generation (1)                [100%] 1 of 1 ✔
[ee/a5bc56] process > decoy_generation (1)                     [100%] 1 of 1 ✔
[b6/694292] process > dia_raw_file_conversion (1)              [100%] 6 of 6 ✔
[27/e264fc] process > dia_spectral_library_search (6)          [100%] 6 of 6 ✔
[c2/205be7] process > dia_search_output_merging (1)            [100%] 1 of 1 ✔
[da/fde7a6] process > global_false_discovery_rate_estimation (1) [100%] 1 of 1 ✔
[9e/cc0474] process > export_of_scoring_results (1)            [100%] 1 of 1 ✔
[10/72ac19] process > chromatogram_indexing (6)                [100%] 6 of 6 ✔
[e1/220cc3] process > chromatogram_alignment (1)               [100%] 1 of 1 ✔
[8f/1d8c62] process > reformatting (1)                         [100%] 1 of 1 ✔
[af/d40fb8] process > statistical_post_processing (1)          [100%] 1 of 1 ✔
[34/a153ac] process > output_visualization (1)                 [100%] 1 of 1 ✔
[2a/9a11d0] process > output_documentation                     [100%] 1 of 1 ✔
-[nf-core/diaproteomics] Pipeline completed successfully-
Completed at: 05-Nov-2020 00:34:00
Duration    : 2h 7m 15s
CPU hours   : 5.4
Succeeded   : 42
```

Figure S2. Benchmarking quantification performance

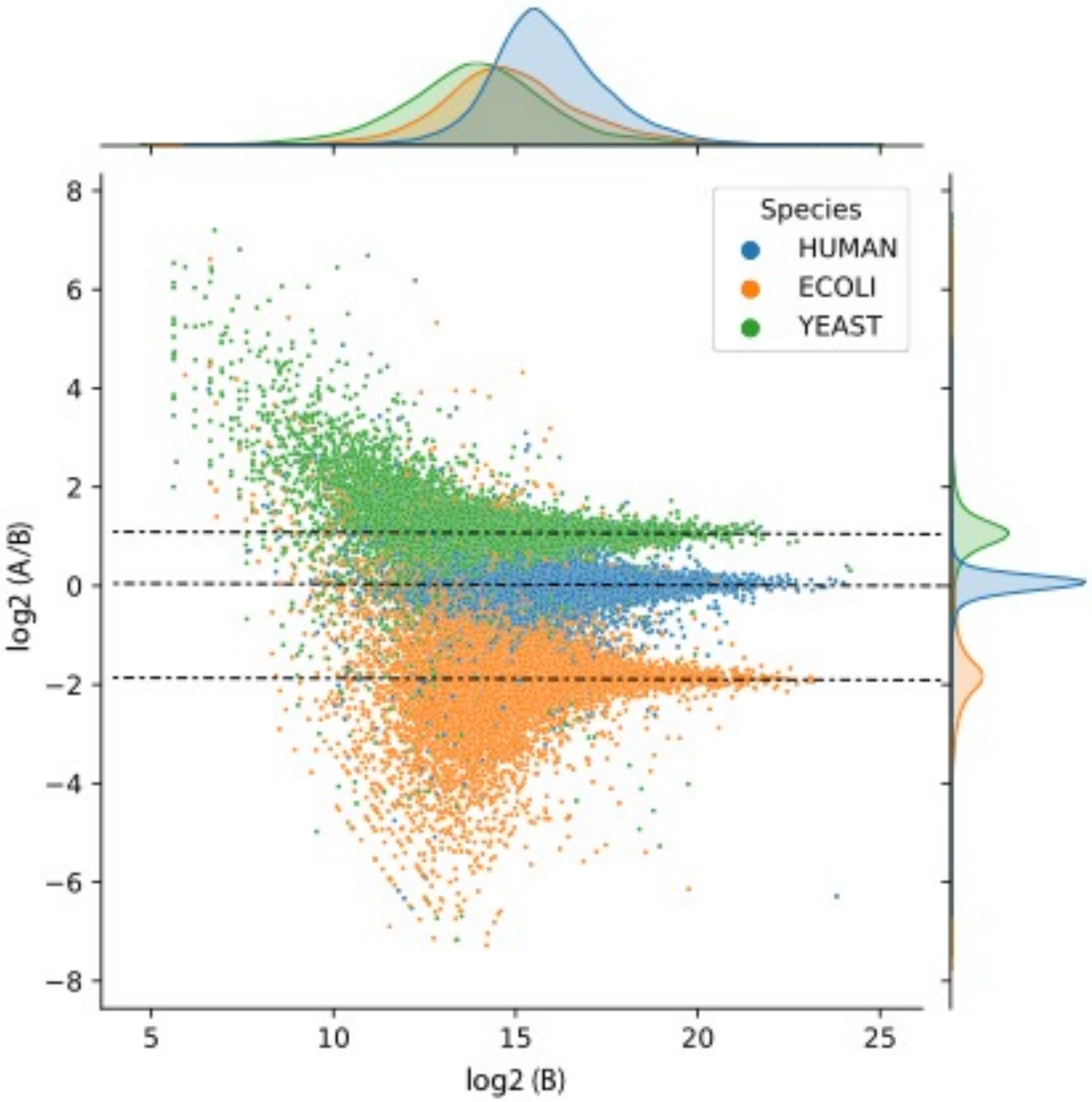


*Reproducing fold changes of human, E. coli and yeast organism mixtures (TripleToF 6600, 64 variable windows) multi-center benchmark study by Navarro P. et al Nature Biotechnology 2017.*

Figure S3. Pairwise RT alignment option for merging multiple spectral libraries

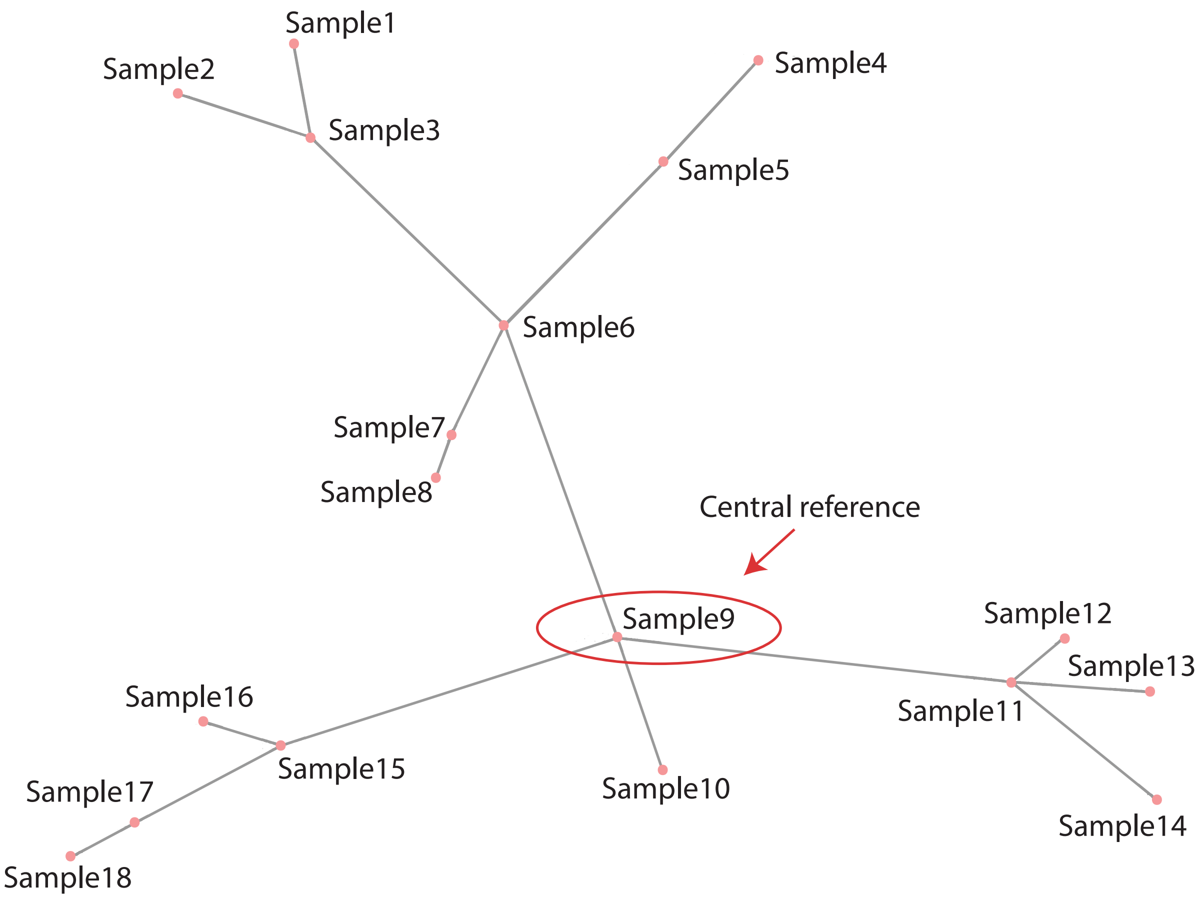


*Example of pairwise RT alignment when merging multiple spectral libraries originating form diverse samples. A minimum spanning tree (MST) based on the peptide share across the samples is constructed and all samples are pairwise aligned along the edges of the MST towards a central reference.*
